# Digital Spatial Profiling of Intraductal Papillary Mucinous Neoplasms: Towards a Molecular Framework for Risk Stratification

**DOI:** 10.1101/2022.09.08.507112

**Authors:** Matthew K. Iyer, Chanjuan Shi, Austin M. Eckhoff, Ashley Fletcher, Daniel P. Nussbaum, Peter J. Allen

## Abstract

The histopathologic heterogeneity of intraductal papillary mucinous neoplasms (IPMN) complicates the prediction of pancreatic ductal adenocarcinoma (PDAC) risk. Intratumoral regions of pancreaticobiliary (PB), intestinal (INT), and gastric foveolar (GF) epithelium may occur with either low-grade dysplasia (LGD) or high-grade dysplasia (HGD). We used digital spatial RNA profiling of dysplastic epithelium (83 regions) from surgically resected IPMN tissues (12 patients) to differentiate subtypes and predict genes associated with malignancy. The expression patterns of PB and GF lesions diverged from INT, suggesting that PB and GF arise from a common lineage. Transcriptional dysregulation within PB lesions mirrored that of PDAC, whereas INT and GF foci did not. Tumor necrosis factor/nuclear factor κB (TNF-NFκB) and cell cycle (cycling-S, cycling-G2/M) programs occurred with relative prominence in PB and INT subtypes, respectively. Taken together, this study delineates markers of high-risk IPMN and insights into malignant progression.

**One Sentence Summary:** Spatial profiling of the intratumoral heterogeneity of IPMN yields markers of high-risk disease and insights into malignant progression.

## INTRODUCTION

Pancreatic ductal adenocarcinoma (PDAC) remains a leading cause of cancer death, predominantly due to the lack of early detection strategies that enable identification of patients at a potentially curable stage*(1)*. Intraductal papillary mucinous neoplasms (IPMN) are cystic lesions of the pancreas that represent a radiographically detectable precursor to pancreatic cancer*(2)*. While the vast majority of IPMN do not progress to malignancy, the ability to accurately differentiate those lesions at low-risk for progression (low-grade dysplasia) from those at high-risk (high-grade dysplasia, early cancer) remains elusive*(3)*. It is widely accepted that operative treatment of high-grade dysplasia (HGD) is appropriate, whereas radiographic surveillance is appropriate for those with low-grade dysplasia (LGD). Consensus guidelines designed to predict risk of high-grade dysplasia through clinical, radiographic, laboratory, endoscopic, and cytologic parameters have an overall accuracy of approximately 60%*(4–6)*. Hence, there has been intense interest in the development of more accurate biomarkers for high-risk IPMN.

Pathologic characterization of IPMN demonstrates multiple dysplastic histologic epithelial subtypes that often coexist within individual specimens. Epithelial subtypes observed histologically include pancreaticobiliary (PB), intestinal (INT), gastric foveolar (GF), and an oncocytic variant. The PB and INT subtypes comprise the overwhelming majority of IPMN with propensity for invasive cancer, and GF represents an indolent subtype associated with favorable prognosis. Studies comparing patient outcomes by subtype suggest that PB histology is more likely to harbor or progress to malignancy, and patients with invasive lesions from PB subtype experience similar outcomes as patients with conventional PDAC*(7–9)*. Genomic interrogation of IPMN has implicated DNA alterations in *KRAS, GNAS*, and *RNF43* as the most prevalent events within neoplasms, but these mutations may coexist and do not reliably associate with histologic subtype or grade of dysplasia*(10–12)*. Mutations such as *TP53, CDKN2A, SMAD4*, and others may be predictors of HGD or invasive carcinoma when present, but their prevalence is relatively low, and the sensitivity is limited. Numerous studies have proposed possible mRNA, microRNA, and protein biomarkers for high-risk IPMN, but to date none has been incorporated into clinical use due to prognostic inaccuracies*(13–18)*.

Intralesional heterogeneity further complicates the discovery of markers for high-risk IPMN. Studies of bulk tissue or cyst fluid convolute the mosaic of epithelial subtypes and grades of dysplasia that occur within individual patients, thereby obfuscating any underlying signal that may be present in disease foci. Microdissection can isolate and compare regions of dysplastic epithelium, but these approaches can be technically challenging and disrupt tissue quality*(19, 20)*. Single-cell RNA-seq of IPMN can characterize unique cell populations within bulk tissues, but do so at the expense of their spatial relationships, and are thus unable to distinguish differences between histologic subtype or pathologic grade*(21)*.

The emergence of multiplex digital spatial profiling addresses the above challenges in deconvoluting heterogeneous tissues to delineate disease phenotypes*(22, 23)*. This technology offers precise comparison of gene expression among user-defined disease regions without the need for cumbersome microdissection. We recently utilized this technology to explore the composition of immune cells within the IPMN tumor microenvironment*(24)*. Here, we report spatial RNA profiling of ductal epithelium across subtypes to determine markers of dysplasia and identify biological processes that associate with malignant progression.

## RESULTS

### Patient characteristics

Digital spatial RNA profiling was performed on FFPE tissue specimens from 12 patients who underwent pancreatectomy for IPMN between 2017 and 2021 (**Fig. 1A**). Clinicopathologic details of the cohort are summarized in **Table 1**. A pancreatic pathologist (CS) characterized the specimens and prepared tissue blocks for profiling according to the following criteria: 1) predominantly INT (N=6) or PB (N=6) histology, 2) presence of at least HGD within the specimen, 3) adequate areas of LGD and HGD within a single tissue block to facilitate a controlled comparison between grades of dysplasia (**Fig. 1B**). Due to the heterogeneity of the tumors, the specimens also contained numerous regions of GF epithelia with uniformly LGD. Invasive carcinoma occurred in four of six PB specimens and none of the INT specimens. To compensate for this potential bias, we prepared slides from areas of the specimen that entirely lacked invasive carcinoma.

**Table 1:** Patient characteristics

|  |  |
| --- | --- |
| Patients | 12 |
| Median Age: years (IQR) | 70 (63-77) |
| Female Sex | 5 (42%) |
| Race |  |
| Caucasian | 8 (67%) |
| Asian | 3 (25%) |
| Black | 0 (0%) |
| Unidentified | 1 (8%) |
| Operation |  |
| Pancreaticoduodenectomy | 9 (75%) |
| Distal Pancreatectomy | 2 (17%) |
| Other | 1 (8%) |
| Median Lesion Size | 5.2cm |
| Invasive Carcinoma Within Lesion | 4 (33%) |
| Cancer Stage | IA (2), IIA (2) |
| Median Follow up (months) | 27 |
| Cancer Recurrence | 2 (50%) |
| Death | 3 (25%) |
| Anatomic Subtype |  |
| Main Duct | 5 (42%) |
| Branch Duct | 3 (25%) |
| Mixed | 4 (33%) |
| Predominant Histologic Subtype |  |
| Intestinal | 6 (50%) |
| Pancreatobiliary | 6 (50%) |
| Gastric Foveolar | 0 (0%) |

**Fig. 1:**
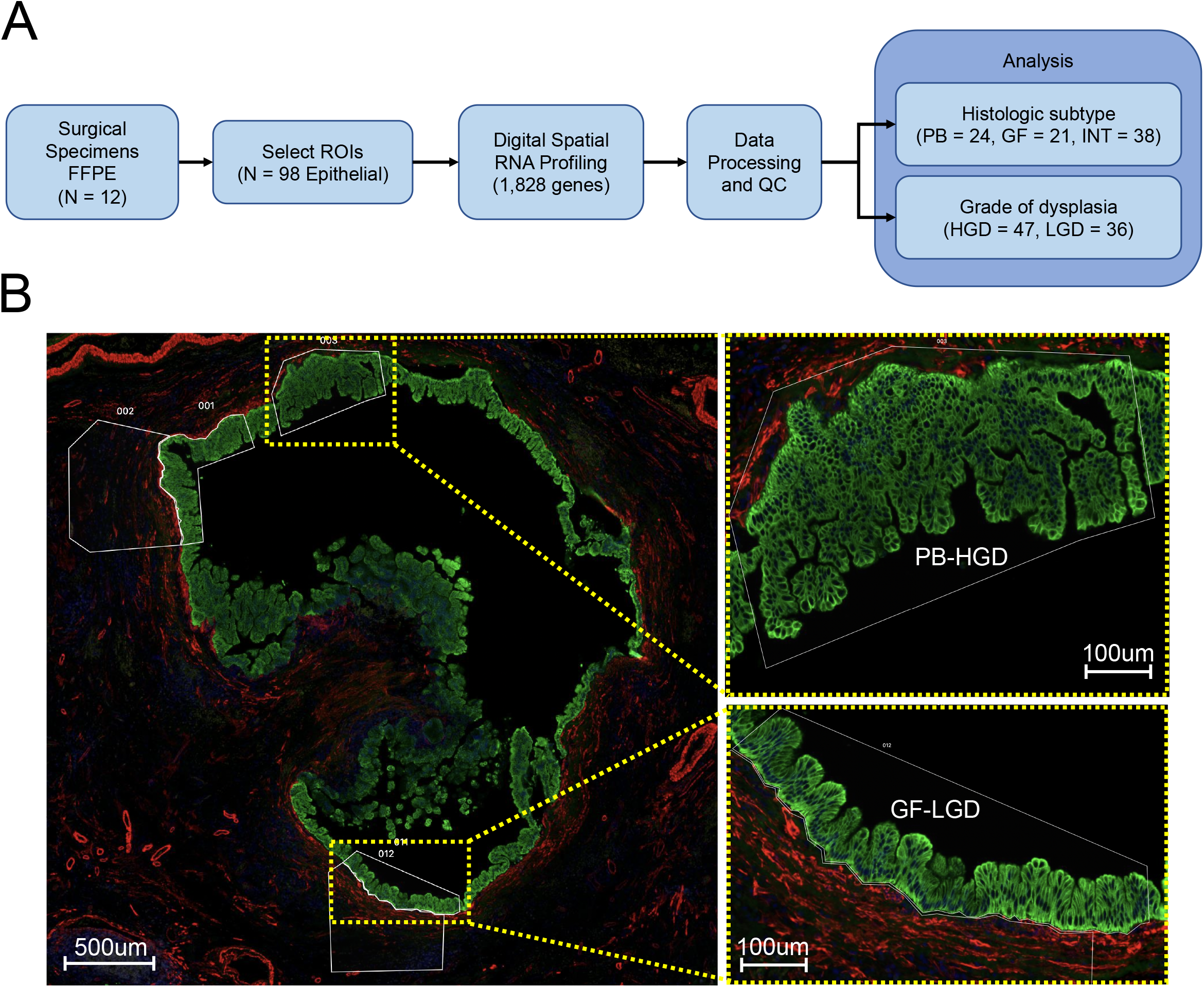
Spatial RNA profiling of IPMN. (**A**) study overview flow diagram, (**B**) a representative high-resolution microscopy image from patient slide 6 depicting regions of HGD (PB) and LGD (GF) cyst epithelium chosen for profiling.

### Targeted spatial transcriptome profiling

We utilized the Nanostring GeoMx Cancer Transcriptome Atlas (CTA) platform to profile a total of 98 epithelial Areas of Interest (AOIs) (50 HGD, 48 LGD) from all slides (6-10 per slide, **Table S1**). The median nuclei count per AOI was 623 (IQR 345-864). High-throughput sequencing yielded a median of 608,651 aligned deduplicated reads per AOI (IQR 261,088 - 1,162,890). Sequencing yield was correlated with AOI nuclei count and surface area, suggesting a strong association with the quantity of *in situ* RNA (**Fig. S1A, B**). Of the 98 AOIs, 83 (84%) met QC filtering criteria (**Fig. S1C**). Epithelial regions with HGD tended to contain greater cellular density than LGD AOIs, leading to increased sequencing counts and signal-to-noise AUC (snAUC) (**Methods, Fig. S1D**). Of the epithelial subtypes, INT harbored the greatest cellular density, followed by PB and finally GF AOIs (**Fig. S1E**). Sequencing yield and snAUC reflect these trends. The normalization strategy to account for this potential bias is discussed below.

A median of 48.3% of probes were expressed above background in each individual AOI (IQR 42.7% - 55.2%, **Table S2**). Gene filtering retained 1,288 / 1,828 (70.4%) genes expressed at detectable levels above background in at least 20% of AOIs (**Fig. S2A, Table S3**). The raw count density distributions of the AOIs indicate that the filtered genes were of low abundance (**Fig. S2B**). Background subtraction followed by quantile normalization was applied to account for differences in sequencing yield, enabling biological comparison of gene expression across AOIs (**Tables S4 and S5**). The normalized count density distributions across individuals AOIs were grossly similar (**Fig. S3A**), and there was no observable bias by either pathologic grade or epithelial histology (**Fig. S3B, C**).

### Divergence of INT from PB and GF subtypes

An unbiased survey of expression patterns was performed using principal component analysis. The AOIs formed two groups that corresponded to histologic subtype, with INT AOIs forming one group and PB-GF (non-INT type) AOIs forming the second (**Fig. 2A**). The separation of AOIs by grade of dysplasia along the first two principal components varied among specimens (**Fig. 2B**). Of the INT patients, the AOIs from slides 4 and 12 grouped tightly together, suggesting homogeny. Two patients (slides 1 and 3) that contained a mixture of GF (LGD) and INT (HGD) epithelium were notable because the GF and INT regions clustered with their respective histologic subtypes, rather than by patient. Notable separation between HGD and LGD AOIs occurred in INT slides 2 and 9. Invasive carcinoma, though excluded from slides prepared for digital spatial profiling, occurred in four specimens with PB histology (slides 5, 6, 8, and 11) (**Fig. 2C**). The HGD AOIs from these four patients grouped closely together, suggesting the existence of a high-risk phenotype. Of the two PB specimens lacking invasive carcinoma, one grouped with the other PB AOIs (slide 7) and other grouped more closely with GF AOIs (slide 10).

**Fig. 2:**
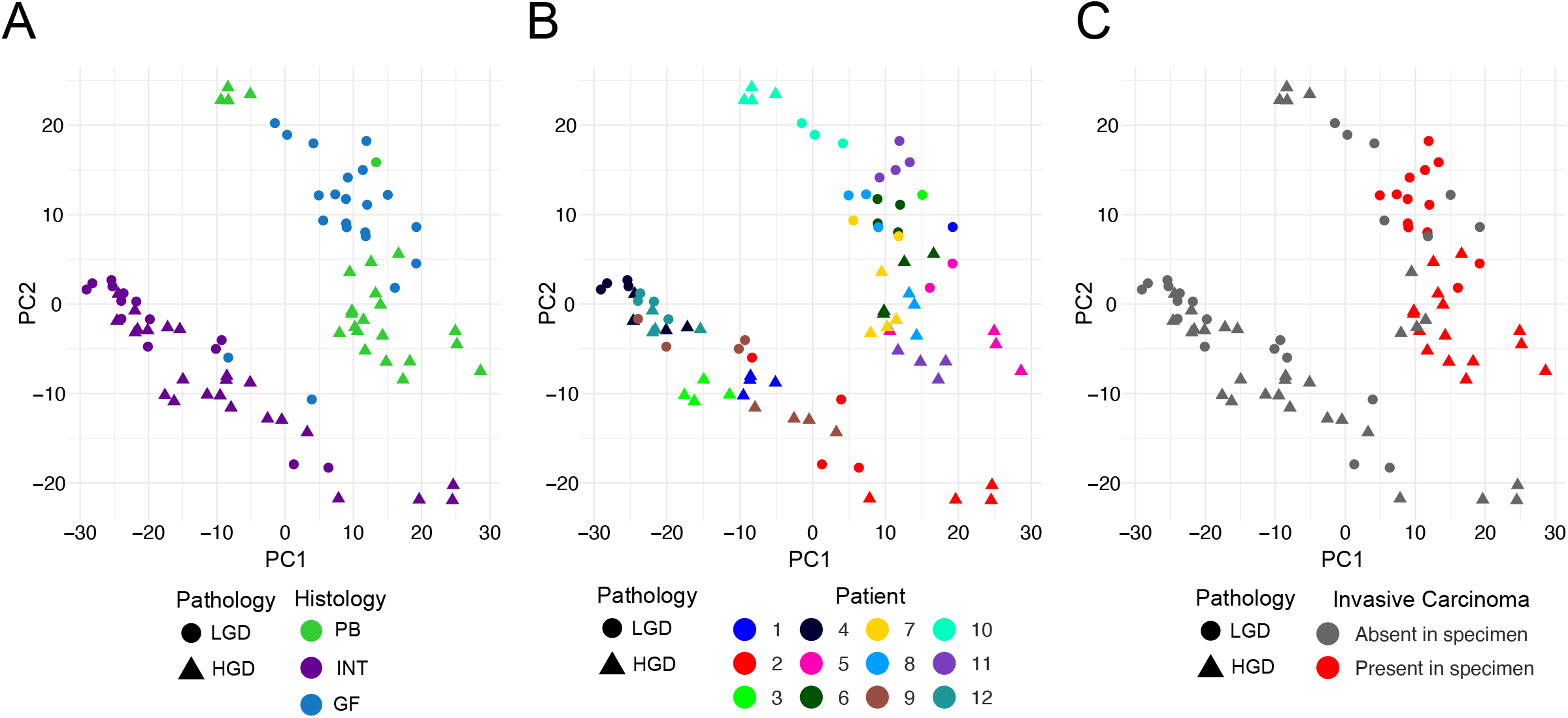
Principal component analysis of normalized spatial RNA profiles. Scatter plot of the first two principal components. Shape of AOIs reflects grade of dysplasia. Panels depict AOIs colored by **(A)** Histologic subtype, **(B)** Patient / Slide ID, and **(C)** Presence of carcinoma elsewhere in the specimen.

### Identification of genes associated with epithelial subtype

DE analysis of PB, INT, and GF AOIs was performed in a pairwise fashion (PB vs. GF, PB vs. INT, INT vs. GF) to discover subtype-specific gene expression (**Table S6**). Intersections of the resulting DE genes formed PB, GF, and INT specific signatures (**Fig. S3**). The union of these sets comprised 127 histologic subtype-specific genes (**Fig. 3A**). Hierarchical clustering of the AOIs produced distinct clusters corresponding to INT, GF, and PB histology. Clustering at the gene level further revealed three clusters denoted C1-INT, C2-PB, and C3-PB-GF (non-INT). The AOIs from slides 2 and 10 did not cluster with their respective epithelial subtypes. Rather, slide 2 clustered with PB AOIs despite having a mix of INT and GF epithelium, and the PB-HGD AOIs from slide 10 clustered with GF (LGD).

**Fig. 3:**
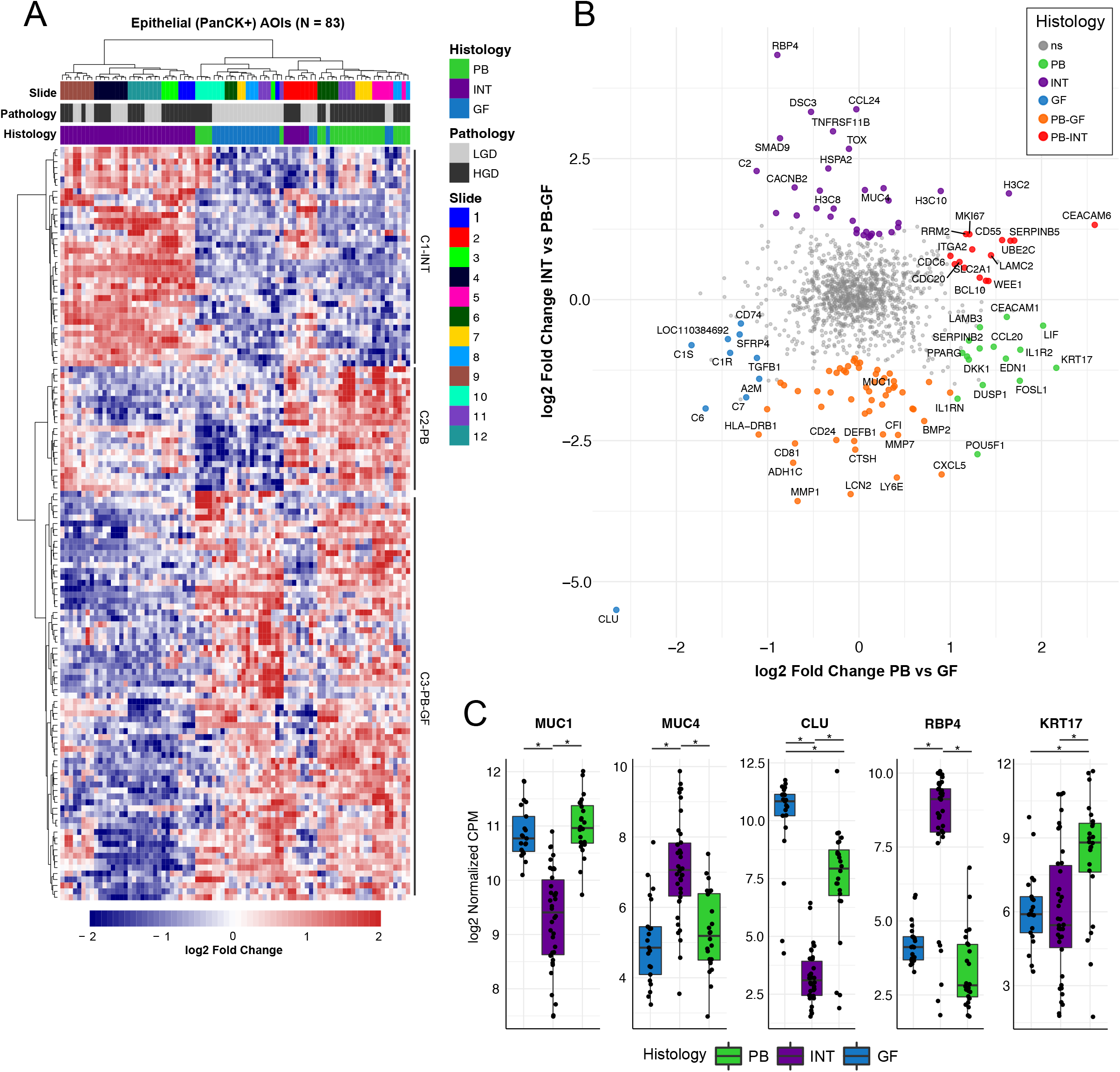
Analysis of epithelial subtypes of IPMN. (**A**) Heatmap plot of differentially expressed genes specific to PB, INT, and GF subtype (absolute log2 fold change > 1, adjusted p-value < 0.05). Columns represent individual AOIs annotated by slide identifier, grade of dysplasia, and epithelial subtype. Rows represent individual genes with expression values scaled by *z-*score. Hierarchical clustering of both columns and rows was performed. (**B**) Scatter plot showing log2 fold change of PB versus GF genes (*x* axis) and INT versus PB-GF (non-intestinal) (*y* axis). Genes that are significantly differentially expressed (absolute log2 fold change > 1, adjusted p-value < 0.05) are colored by the histologic subtype that they represent. The top 10 differentially expressed genes from each analysis are labeled. The widely used marker gene *MUC1*, which was overexpressed in PB-GF relative to INT, is labeled for reference. (**C**) Boxplots showing the log-normalized gene expression of subtype marker genes *MUC1, MUC4, CLU, RBP4*, and *KRT17*.

When describing histologic subtype marker genes across the three subtypes, the presence of known IPMN marker genes served as external validation for the DE results. Mucin genes, including *MUC1, MUC2, MUC4, MUC6*, and *MUC5AC*, have been proposed to discriminate between subtypes*(14, 15)*, and the NanoString CTA probe set contained two of these mucin genes, *MUC1* and *MUC4*. Consistent with prior reports, *MUC1* was overexpressed in PB-GF (non-INT) AOIs, and *MUC4* was overexpressed in INT AOIs*(14, 15, 25–27)* (**Fig. 3B and 3C**). At the RNA level, neither *MUC1* nor *MUC4* were found to be significantly altered in PB relative to GF AOIs. Notably, several genes outperformed *MUC1* and *MUC4* as IPMN subtype marker genes, including Clusterin (*CLU*), Retinol binding protein 4 (*RBP4*), and Keratin 17 (*KRT17*), which showed marked overexpression in GF, INT, and PB AOIs, respectively (**Fig. 3C**).

### Identification of genes associated with high-risk IPMN

DE analysis of HGD versus LGD AOIs yielded 38 significant genes (30 overexpressed, 8 underexpressed), **Fig. 4A, Table S6**). Semi-supervised clustering using these genes produced two distinct clusters of AOIs: 1) a “high-risk” cluster containing 33 / 83 AOIs (∼40%) from 5/6 PB slides (5, 6, 7, 8, and 11) and 3/6 INT slides (1, 2, and 9), and 2) a “low-risk” cluster containing all GF AOIs and HGD AOIs from slides 3, 4, 10, and 12.

**Fig. 4:**
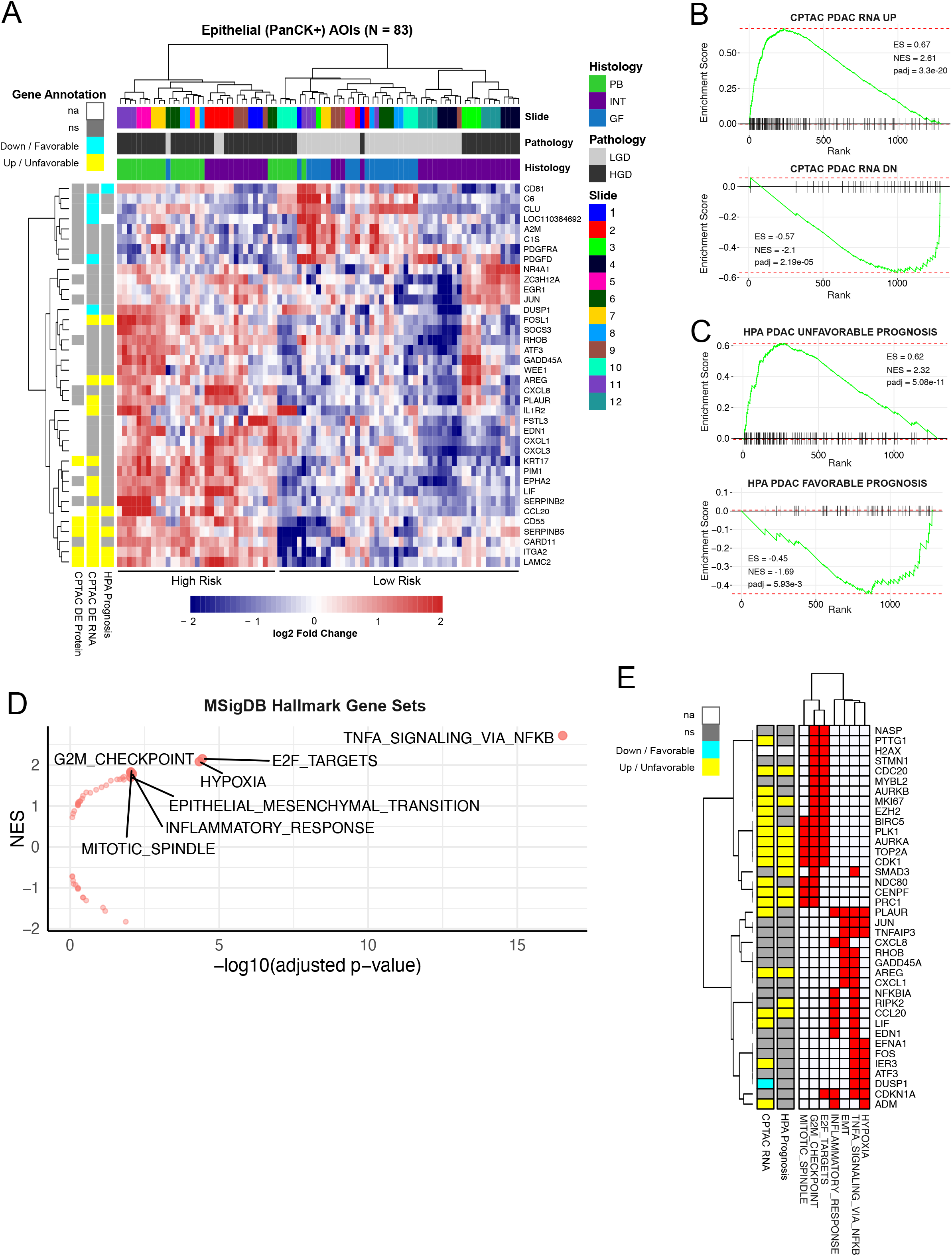
Analysis by grade of dysplasia (HGD versus LGD). (**A)** Heatmap plot of differentially expressed genes in AOIs representing HGD versus LGD (absolute log2 fold change > 1, adjusted p-value < 0.05). Gene expression values are scaled by *z*-score. AOIs (columns) are annotated by slide identifier, grade of dysplasia, and epithelial subtype. High risk and low risk AOI clusters are annotated below. Genes (rows) are further annotated by established markers PDAC, including datasets from the Human Protein Atlas (HPA Prognosis) and CPTAC consortium analysis (CPTAC DE RNA, CPTAC DE Protein). **(B-C)** GSEA enrichment plots using external PDAC gene sets comparing AOIs with HGD compared to LGD, ranked by log fold change. **(D)** Volcano plot of MSigDB Hallmark gene sets associated with HGD compared to LGD, with statistical significance plotted on the *x* axis and normalized enrichment score plotted on the *y* axis. Significantly enriched gene sets (adjusted p-value < 0.01) are shown with text labels. **(E)** Heatmap showing genes (rows) associated with two or more enriched Hallmark gene sets (columns). Rows are also annotated with evidence of dysregulation (CPTAC RNA) or prognostic relevance (HPA Prognosis) in PDAC.

Given that IPMN with HGD represents PDAC *in situ*, we postulated that gene expression changes essential to the progression to HGD should be retained in PDAC*(8)*. To investigate this rationale we compared DE genes in HGD versus LGD IPMN with PDAC gene sets curated from external sources: 1) Clinical Proteomic Tumor Analysis Consortium (CPTAC) RNA-Seq and proteomics analysis of pancreatic cancer versus normal adjacent tissues, 2) the Human Protein Atlas (HPA) analysis of The Cancer Genome Atlas (TCGA) RNA-Seq data predicting prognostic genes in pancreatic cancer, 3) Mao *et al*. analysis of bulk RNA-Seq data from PDAC versus adjacent benign tissue, and 4) Grutzmann *et al*. meta-analysis of pancreatic cancer microarray experiments (see Materials and Methods) *(28–31)*. The analyses from CPTAC and HPA (TCGA) were considered more robust as these were generated by large consortia using standardized protocols. Of the thirty genes overexpressed in HGD versus LGD IPMN, six were overexpressed in PDAC (CPTAC RNA-Seq) and associated with unfavorable prognosis (HPA): *FOSL1, AREG, CCL20, SERPINB5, ITGA2, LAMC2* (**Fig. S4**). Additional genes found to be overexpressed in both IPMN and PDAC but not associated with prognosis included *PLAUR, IL1R2, KRT17, EPHA2, LIF, CD55*, and *CARD11*.

Gene Set Enrichment Analysis (GSEA) demonstrated significant enrichment (adjusted p-value < 0.05) against all curated gene sets (**Fig. S5, Table S7**). Statistical significance largely depended on gene set size, with downregulated genes exhibiting weaker enrichment than upregulated genes. The most significant enrichment was observed with the RNA-Seq analysis from CPTAC (**Fig. 4B**) and the HPA analysis of prognosis (**Fig. 4C**).

To search for molecular mechanisms underlying progression of IPMN, we performed exploratory GSEA against the MSigDB Hallmark gene set collection (**Fig. 4D, Table S8**)*(32)*. Genes were ranked by log fold change in HGD versus LGD IPMN. This resulted in 7/50 significantly enriched gene sets (padj < 0.01). The gene set with the highest enrichment score represented genes regulated by NF-κB in response to tumor necrosis factor (TNF) signaling (TNF-NFκB). We interpreted this result as supportive of the known link between inflammatory signaling and progression in IPMN*(33–36)*. An additional 3/7 enriched gene sets pertained to cell proliferation. Leading edge analysis of the genes enriched in two or more genes sets yielded two unique clusters (**Fig. 4E**). The first cluster included *MKI67* and other genes associated with cell proliferation and division, and the second involved genes associated with inflammatory signaling, hypoxia, and epithelial-to-mesenchymal transition (EMT).

### Genes that differentiate high-grade PB from INT IPMN

We next investigated neoplastic progression in PB and INT-predominant tumors. IPMN specimens were partitioned by their predominant histology subtype across the entire specimen. Accordingly, the AOIs formed four subgroups: PB-HGD (N = 23), PB-LGD (N = 18), INT-HGD (N = 24), and INT-LGD (N = 23). The PB-LGD subgroup included GF AOIs (N = 17) and a single low-grade PB AOI. There were 50 DE genes (34 overexpressed, 16 underexpressed) specific to the PB-HGD versus PB-LGD analysis, and 6 genes (5 overexpressed, 1 underexpressed) specific to the INT-HGD versus INT-LGD analysis (**Fig. 5A, Fig. S6, Table S6**). Carcinoembryonic Antigen-Related Cellular Adhesion Molecule 6 (*CEACAM6*), which has been reported as a possible biomarker in IPMN and cholangiocarcinoma, was highly upregulated in PB but not INT IPMN*(21, 37)* (**Fig. 5B**). Additionally, Laminin subunit beta-3 (*LAMB3*) and POU Class 5 Homeobox 1 (*POU5F1*) were specific to PB-HGD but not INT-HGD AOIs. Early growth response-1 (*EGR1)* was more specific to INT, though met significance thresholds in the grouped analysis of HGD versus LGD dysplasia.

**Fig. 5:**
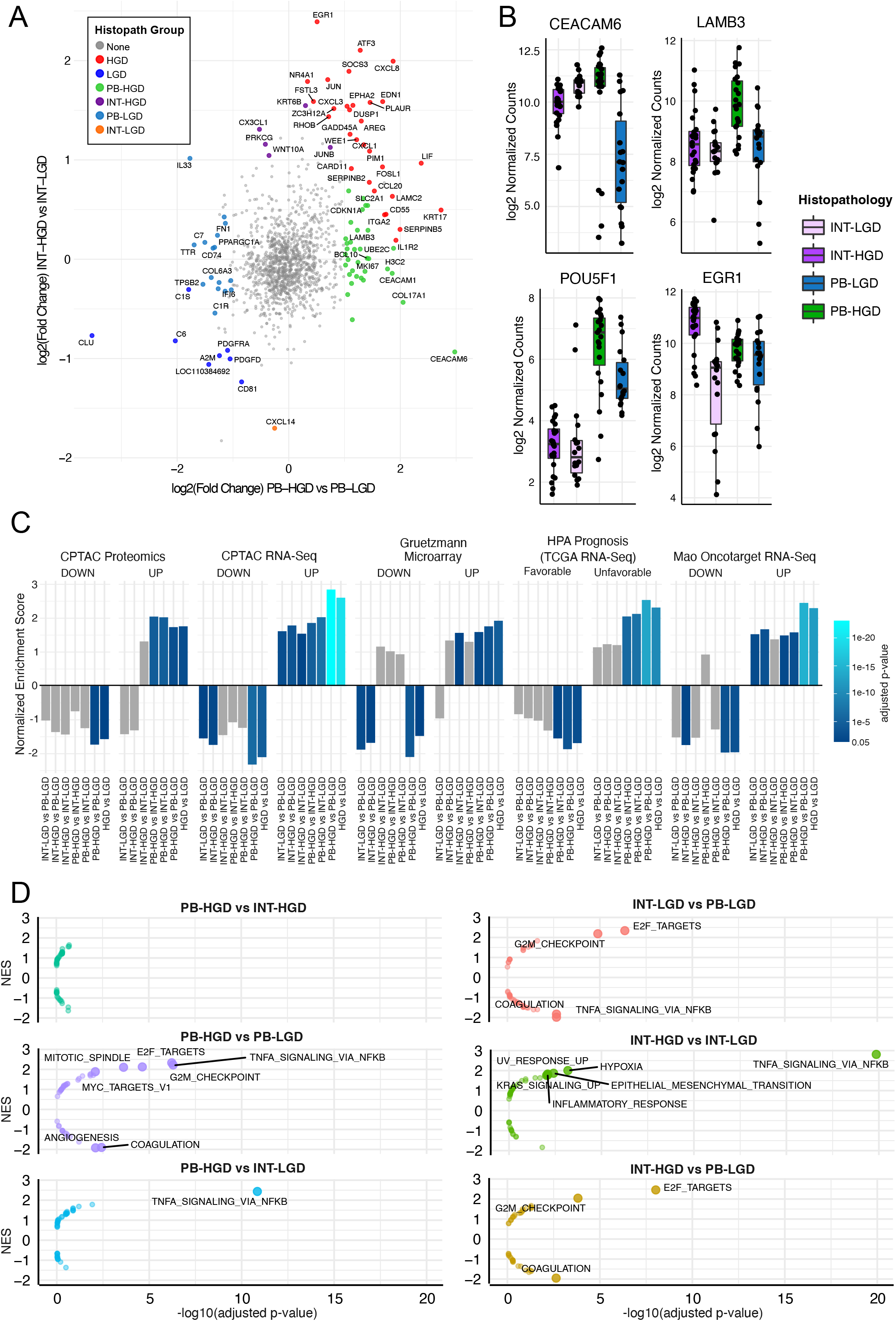
Comparison of epithelial subtypes by grade of dysplasia. (**A**) Scatter plot showing log2 fold change of PB-HGD versus PB-LGD (*x* axis) and INT-HGD versus INT-LGD (*y* axis). Significant DE genes are colored by histopathologic subgroup. Top 10 DE genes from each analysis are labeled. (**B**) Boxplots showing the log-normalized gene expression of histopathologic marker genes. (**C**) Bar plots depicting GSEA normalized enrichment score obtained from testing of histopathologic groups against several external PDAC gene sets. Bar colors depict the adjusted p-value of each test. Gray bars are not statistically significant (padj > 0.05). (**D**) Volcano plots of MSigDB Hallmark gene sets associated with each histopathologic subgroup (relative to its counterparts), with statistical significance plotted on the *x* axis and normalized enrichment score plotted on the *y* axis. Significant enrichment results (adjusted p-value < 0.01) are shown with text labels.

### Association of IPMN subtypes with PDAC

Subgroup DE analysis was performed on pairwise combinations of histopathologic groups (6 comparisons) and the resulting ranked gene lists were tested against the curated PDAC gene sets (**Table S7**). The PB-HGD versus PB-LGD analysis had the greatest absolute normalized enrichment score (NES) in 8/10 gene sets and outperformed analyses involving INT-HGD regions for every gene set (**Fig. 5C**). Of particular significance was the stark difference in NES for the HPA Prognosis gene sets. All comparisons involving PB-HGD AOIs were highly enriched, whereas none of the INT-HGD comparisons showed significant enrichment. This suggests that PB-HGD IPMN most closely resembles invasive carcinoma and may represent a direct precursor to malignancy in the dysplastic progression of IPMN.

### Upregulation of inflammatory signaling and cell proliferation during dysplastic progression

Exploratory GSEA against MSigDB Hallmark gene sets was replicated for each of the 6 pairs of histopathologic groups (**Fig. 5D, Table S8**). Cell proliferation programs (S and G2/M phases) were upregulated in INT-LGD relative to PB-LGD (GF) AOIs. This was concordant with the greater cell density observed within INT regions on microscopic examination and reflected in the initial QC analysis (**Fig. S1E, Fig. S8**). By contrast, the TNF-NFκB transcriptional program was upregulated in PB-LGD relative to INT-LGD AOIs. Both PB-HGD and INT-HGD regions overexpressed both TNF-NFκB and proliferation pathways relative to their LGD counterparts. Taken together, these results suggest that INT may arise as a primarily proliferative lesion before acquiring the TNF-NFκB program and progressing to HGD. By contrast, PB-LGD (GF) may arise in the setting of inflammatory signaling and acquire the capacity to proliferate during progression to HGD. No significant hallmark pathways distinguished PB-HGD from INT-HGD, suggesting that transcriptional programs not encompassed by MSigDB Hallmark gene sets must account for the marked differences between PB-HGD and INT-HGD.

### Unsupervised network analysis to delineate gene clusters associated with progression to carcinoma

To determine whether gene expression patterns could infer biological pathway activity and cancer risk without reliance on pathological annotation, we performed co-expression network analysis (Materials and Methods). This resulted in a network containing 300 genes connected by 791 edges (**Fig. 6A, Tables S9 and S10**). Unsupervised clustering partitioned the network into 29 clusters of co-expressed genes. We discarded clusters with <10 genes (23 / 29) for which enrichment testing would be underpowered, leaving 6 communities for further investigation.

**Fig. 6:**
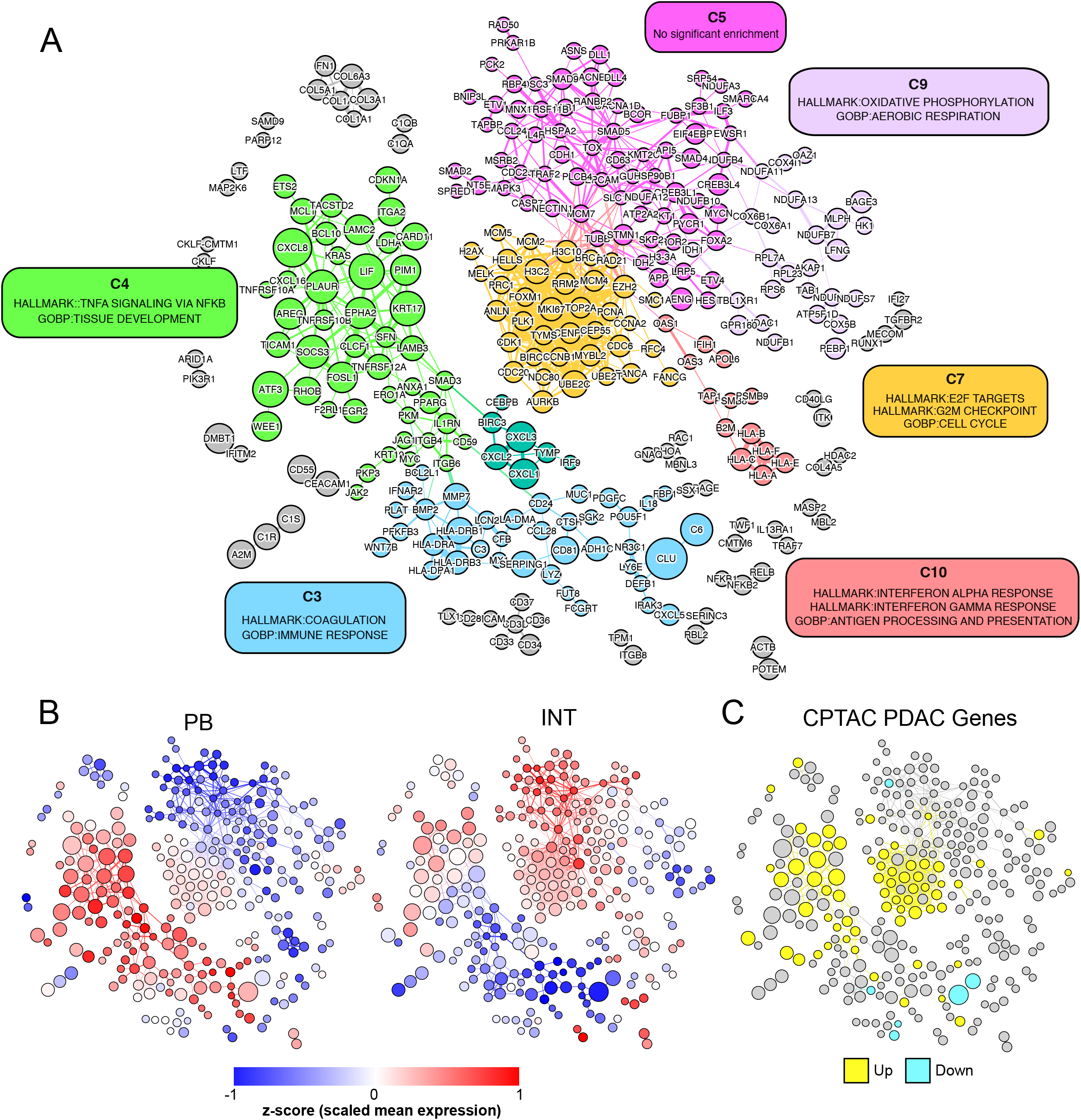
Co-expression network analysis. Gene co-expression network was produced by correlation analysis of all epithelial AOIs, where nodes represent individual genes and edges connect highly correlated gene pairs. Node size reflects the fold change in expression between HGD and LGD AOIs.(**A**) Unsupervised clustering of network with color-coded clusters. Clusters with >10 genes were tested for enrichment against MSigDB Hallmark gene sets, GO:BP gene sets, and PDAC gene sets. Cluster annotation based on the most significantly enriched gene sets are shown in colored boxes. (**B**) Overlay showing standardized mean gene expression (z-score) across PB-HGD AOIs (left) and INT-HGD AOIs (right). (**C**) Overlay showing overexpressed (yellow) and underexpressed (cyan) genes from the CPTAC PDAC RNA-Seq dataset.

Hypergeometric enrichment testing was used to associate clusters with gene sets from the MSigDB Hallmark database as well as Gene Ontology Biological Processes (GO:BP, **Table S11**)*(38)*. After consolidating redundant biological processes, we selected the most statistically significant gene sets to represent each cluster (**Fig. 6A**). We then projected the standardized mean gene expression across PB-HGD and INT-HGD AOIs, respectively, onto the network using color gradient overlays (**Fig. 6B**). To relate network clusters with PDAC we performed enrichment testing against PDAC gene sets and visualized the CPTAC RNA-Seq DE genes as a third network overlay (**Fig. 6C**).

Clusters C4 and C7 warranted consideration as contributors to malignant progression based on significant enrichment for CPTAC RNA-Seq PDAC genes (C4 22 / 46 genes, adjusted p-value = 3.7e-6, C7 25 / 37 genes, adjusted p-value = 8.3e-12). Cluster C4 was significantly enriched for TNF-NFκB signaling (17 / 46 genes, adjusted p-value = 4.4e-6) and contained 15 / 30 genes overexpressed in HGD versus LGD IPMN. Genes within C4 tended to be expressed at higher levels in PB relative to INT IPMN. Accordingly, C4 harbored 15 genes overexpressed in PB-HGD versus PB-LGD AOIs, compared to just 6 genes overexpressed in INT-HGD versus INT-LGD AOIs. Cluster C7 was significantly enriched for cell cycle genes (28 / 37 genes, adjusted p-value = 2.1e-18). Genes within C7 tended to be expressed at higher levels in INT relative to PB IPMN, and contained more genes upregulated in INT versus GF AOIs (22 / 37 genes) than PB versus GF AOIs (6 / 37 genes). In contrast to C4, C7 contained none of the DE genes in HGD versus LGD IPMN. Taken together, these results implicate C4 (TNF-NFκB) as a prominent transcriptional program in PB IPMN and C7 (cell cycle) as a prominent program in INT IPMN.

## DISCUSSION

Management of patients with IPMN presents an opportunity to prevent pancreatic cancer, however current management strategies are limited in their ability to provide accurate recommendations because our ability to predict timing of progression is limited. Cytologic or pathologic confirmation of disease subtype and/or grade of dysplasia is difficult without operative resection, and resection is associated with substantial morbidity and even mortality. Asymptomatic patients who present without high-risk radiographic stigmata of carcinoma represent a clinical conundrum for which no accurate diagnostic modalities currently exist. For this patient population, the concept of a prognostic molecular assay holds great promise, but despite considerable investigation no such assay has gained clinical traction.

Available retrospective evidence suggests an association between histopathology and clinical outcomes in patients with IPMN. Specifically, PB histologic subtype portends poor prognosis relative to INT, GF appears to represent an indolent entity, and the presence of HGD forecasts the development of invasive cancer*(7, 39–42)*. Numerous efforts have failed to translate these retrospective pathologic observations into predictive biomarkers for technical and disease-related reasons. The major caveat with “bulk” studies of IPMN tissue rests in the assignment of a single grade and subtype annotation to a specimen, effectively homogenizing the disease morphologies and degrees of dysplasia that occur within the affected pancreas. In an unsupervised clustering analysis of single cell RNA-Seq data from patients with IPMN, Bernard *et al*. found subpopulations of cells from tissues designated LGD within clusters of HGD and carcinoma cells, corroborating histopathologic evidence that IPMN harbor a mixture of cells along a dysplastic spectrum*(21)*. Ultimately, our understanding of the neoplastic progression of IPMN hinges upon the ability to characterize these tissues in a more granular fashion.

Until now, attempts to isolate neoplastic ductal epithelium required technically challenging tissue handling. Jury *et al*. combined laser capture microdissection (LCM) with microarray technology to study gene expression changes associated with IPMN progression*(20)*. The investigators reported markers of pancreatic islet cells, including the hormones insulin, glucagon, and somatostatin, among the most significantly differentially expressed genes in their dataset, raising doubt that the microdissection procedure precisely isolated neoplastic epithelium. Sato *et al*. employed selective microdissection paired with microarrays to compare gene expression between normal ductal epithelia, non-invasive IPMN, and invasive IPMN*(19)*. The authors reported several genes also found by our study, including Serpin family B member 5 (*SERPINB5*), CD55 Molecule (Cromer Blood Group) (*CD55*), and Integrin subunit alpha 2 (*ITGA2*). However, the study exclusively examined regions of carcinoma *in situ* (HGD) and lacked classification by epithelial subtype, limiting its clinical translational potential. As an alternative to tissue isolation, IPMN cyst fluid, rich in DNA, RNA, and protein, can be obtained with minimally invasive fine needle aspiration. However, the acquired material constitutes a convolution of secreted molecules and sloughed off debris from normal and neoplastic pancreatic tissue. Therefore, biomarker identification from either cyst fluid or bulk tissue present similar challenges.

In the current study, we leveraged digital spatial RNA profiling using a targeted gene panel to characterize precise regions of IPMN histopathology across tissue slides and produce robust gene expression patterns by epithelial subtype and grade. Unsupervised dimensionality reduction analysis demonstrated distinct groups of INT and non-INT (PB-GF) AOIs. Importantly, INT and GF AOIs clustered apart even when derived from the same patient and in the same tissue section. By contrast, the intimate association of PB and GF AOIs suggests that GF and PB share a common neoplastic cell lineage distinct from INT lesions. Given that GF regions are almost universally deemed low-grade, we suggest that GF can essentially be considered a precursor to PB epithelium.

Currently, mucin genes serve as a generally accepted differential marker of epithelial subtype. The two mucin genes (*MUC1* and *MUC4*) included in the CTA panel showed significant association with PB-GF and INT subtypes, respectively. However, despite the intended use of *MUC1* as a specific marker of PB IPMN, it did not discriminate between PB and GF AOIs in this dataset. Rather, clusterin (*CLU*), retinol binding protein 4 (*RBP4*), keratin 17 (*KRT17*), and other candidate marker genes nominated by this analysis displayed superior potential to classify epithelial subtypes.

Comparison of gene expression by grade of dysplasia identified gene expression alterations that mirrored those reported in PDAC. Evaluation of gene expression enrichment against independent datasets also served as validation of the spatial profiling platform and our data analysis approach. Clustering the AOIs on the set of 38 genes dysregulated in HGD versus LGD partitioned the specimens into high risk and low risk groups. Eight of the twelve slides, including the four slides from patients with invasive carcinoma, harbored one or more high-risk AOIs. We envision that a gene expression classifier derived from this gene signature could serve as a risk stratification tool for patients with IPMN.

This study also corroborates evidence for individual candidate biomarkers of high-risk IPMN, including *CD55*, laminin subunit gamma 2 (*LAMC2*), amphiregulin (AREG), and others. As noted above, *CD55* was previously reported as a biomarker for IPMN by gene expression microarray experiments, and was found to be associated with disease progression in a proteomic profiling study of IPMN cyst fluid*(19, 43)*. An enzyme-linked immunosorbent assay (ELISA) of *LAMC2* in plasma of PDAC patients augmented the accuracy of the widely used PDAC biomarker CA-19-9*(44). LAMC2* has also been detected in pancreatic duct fluid exosomes in patients with IPMN and PDAC*(45). AREG* was found to be predictive of high-risk IPMN in a serum biomarker panel based on antibody microarray technology and has also been detected in pancreatic cyst fluid*(46, 47)*. Validation studies using these and other candidate genes in pancreatic cyst fluid are warranted.

To explore the biological underpinnings of IPMN we performed a combination of supervised and unsupervised analyses. The supervised analysis leveraged annotation of AOIs by a pancreatic pathologist, differential expression testing, and GSEA to discover biological associations. The unsupervised analysis utilized gene expression correlation information to construct a co-expression network and a community detection algorithm to partition the network into clusters of highly correlated genes. Clusters were then assessed for biological significance through hypergeometric enrichment testing. Ultimately, the two analysis approaches led to similar conclusions. Two transcriptional programs appear to be driving neoplastic progression in IPMN: inflammatory signaling (TNF-NFκB) and cell proliferation (S and G2/M phases). Activation of cell proliferation was more prominent in INT relative to PB lesions, whereas inflammatory signaling was more pronounced in PB than INT. Malignant potential, assessed by enrichment of gene alterations shared with PDAC datasets, was predominantly associated with PB epithelium. The unsupervised network analysis corroborated these findings, yielding a 46-gene cluster associated with the constellation of PB epithelium, inflammatory signaling, and PDAC genes, without *a priori* knowledge of pathologic annotations. Measurement of the transcriptional activity of this gene signature could transcend the histopathologic designations that currently serve as surrogate measures of malignancy risk.

This study has several important limitations. First, the CTA probe panel used in this study measures only ∼10% of human protein coding genes. A targeted panel reduces the power of GSEA and hypergeometric testing alike to detect significant biological pathway enrichment. Genes not measured by the CTA panel may outperform the marker genes nominated by this study or provide evidence of other transcriptional programs with relevance in IPMN. Incorporation of a more comprehensive panel will be important in future studies.

Second, our modest cohort size of twelve specimens may not embody the breadth of disease biology across the common epithelial subtypes (PB, GF, INT) and omits rare entities such as oncocytic IPMN. In addition, the co-occurrence of invasive carcinoma in the majority of the PB cohort and none of the INT cohort confounds our comparison of the two subtypes. Our attempt to mitigate this confounder by requiring all tissue slides to be devoid of invasive carcinoma may or may not be compensatory. Certainly, the available clinical outcomes data support our findings: patients with invasive carcinoma derived from INT fare far better than patients with PB-derived carcinomas*(7)*. Expanded profiling that includes specimens with and without invasive carcinoma from both subtypes will be needed to fully resolve this issue.

A third and related limitation of the cohort design was the requirement that every specimen in the study possess regions of HGD. It is conceivable that regions of LGD from specimens lacking HGD/invasive carcinoma could be different from areas of LGD found in conjunction with HGD/invasive carcinoma. The idea that high-risk IPMN could be detected prior to the development of HGD would certainly alter our clinical approach to the disease. Expanded spatial profiling that includes regions of normal ductal epithelium and invasive carcinoma could address this intriguing possibility.

In summary, our findings offer several refinements to our understanding of IPMN. First, GF epithelium likely represents a precursor to PB rather than a common progenitor to either PB or INT. Second, the activation of inflammatory signaling associates with high-risk IPMN and occurs predominantly in PB lesions. This finding lends credence to ongoing clinical trials of anti-inflammatory therapies in the prevention of IPMN progression. Finally, the incorporation of subtype-specific and high-risk marker genes nominated by this study may facilitate the development of an accurate risk stratification assay in IPMN.

## MATERIALS AND METHODS

### Patient recruitment

Archival biospecimens from patients who had undergone pancreatic resection for IPMN at Duke University Hospital System (DUHS) between 2017 and 2021 were considered for spatial RNA profiling. The Institutional Review Board (IRB) approved the use of de-identified patient specimens for retrospective molecular profiling. Informed consent was not required due to the retrospective nature of the study with minimal risk and de-identification of the specimens. Clinicopathological data was collected by study coordinators and securely stored in a RedCap database. Archived FFPE specimens were procured and reviewed by a board-certified pathologist specializing in pancreatic pathology to confirm diagnosis (CS). Specimen blocks were cut into 5 μm thick serial sections. One section was stained with hematoxylin and eosin (H&E) and imaged using a Nikon TE2000-E microscope for pathology review. Sections containing regions of both LGD and HGD were selected for spatial transcriptomics and mounted on a positively charged slide for this application.

### Spatial RNA profiling

Digital spatial RNA profiling was conducted using the NanoString GeoMx Digital Spatial Profiler (DSP)*(22)*. Our pathologist (CS) selected Regions of Interest (ROIs) annotated by histologic subtype (PB, GF, INT) and grade of dysplasia (LGD or HGD). We selected ROIs that encompassed the spectrum of subtype-grade combinations present on each slide and included multiple biological replicates of each combination. The GeoMx DSP imposes a maximum ROI diameter of 700um. Individual ROIs were drawn to maximize the number of epithelial cells contained while adhering to the size constraint. Segmental profiling of individual cell populations within each ROI was performed by staining the tissues with fluorescently conjugated antibodies: CD45 for immune cells, smooth muscle actin (SMA) for stromal fibroblasts, and anti-pan-cytokeratin (PanCK) for epithelial cells. DSP tissue slides were incubated with the fluorescently conjugated antibodies to CD45, PanCK, and SMA along with a cocktail of photo-cleavable-oligonucleotide probes from the GeoMx CTA kit. Segmentation thus produced multiple Areas of Interest (AOIs) from each ROI. Libraries were prepared according to the NanoString GeoMx Library Preparation Manual and pooled to equimolar concentration. RNA was sequenced under standard conditions on an Illumina Novaseq 6000 to a depth of 30 read-pairs/μm^2^.

### Quality control and normalization

UV-cleaved barcode sequencing reads were processed by the GeoMx DSP Analysis Server. Processing steps included read trimming, alignment, and deduplication. A tabulated matrix of probe counts for each AOI was exported from the DSP server and subsequently analyzed using R version 4.2.0.

Quality control (QC) metrics for each AOI were computed, including the receiver operating characteristic (ROC) curve and the associated area under the curve (AUC) metric of gene probes (N = 8,584) versus negative probes (N = 75)*(48)*. This was used to denote the signal-to-noise AUC (snAUC). A limit of detection (LOD) for each AOI was set to the 90^th^ percentile count of negative probes. A probe count below this LOD was considered undetectable. The following quality control criteria were used to filter AOIs: 1) >100,000 aligned deduplicated total counts and 2) snAUC >0.65. AOIs that did not meet these QC criteria were removed.

The CTA kit features multiple probes per gene (8,584 independent probes targeting 1,829 unique gene identifiers). Probes with undetectable expression in >80% of AOIs were removed, and the geometric mean of representative probe counts was computed to produce a single expression value for each gene. Raw gene counts were normalized in two steps to account for differences in background levels and sequencing output across AOIs. The geometric mean of negative probe counts, a measure of background level, was subtracted from each AOI. Background-subtracted counts were scaled by total library size and subjected to quantile normalization*(49)*.

### Data analysis

Analysis of filtered, normalized gene expression data was performed in the R language with Bioconductor*(50)*. Dimensionality reduction analysis was performed with principal components analysis (PCA). Differential expression (DE) analysis was conducted using the *limma* package*(51)*. Criteria for calling DE genes included absolute log2 fold change > 1.0 and adjusted p-value < 0.05. Heatmap plots were generated using the *pheatmap* R package with row and column clustering using the “ward.D2” method*(52)*. Gene Set Enrichment Analysis (GSEA) was conducted using the Bioconductor package *fgsea(53–55)*. Genes were ranked by their log2 fold change in each DE analysis (e.g., HGD vs. LGD). Gene sets from MSigDB version 7.5.1. were obtained from the *msgidbr* package *(55, 56)*.

### Curation of external datasets

External RNA-Seq analysis results from the National Cancer Institute’s Clinical Proteomic Tumor Analysis Consortium (CPTAC) proteogenomic characterization of pancreatic adenocarcinoma were obtained from Supplementary Table S3*(28)*. Genes were merged by official gene symbol. Genes with an absolute log2 fold change ≥ 1.0 and adjusted p-value < 0.05 were considered differentially expressed and included in gene sets. RNA-Seq analysis from Mao *et al*. comparing PDAC versus matched normal pancreas was curated into gene sets (log2 fold change ≥ 1.0, adjusted p-value < 0.01)*(29)*. Prognostic pathology information from the Human Protein Atlas (HPA) resource was downloaded from the HPA website (https://www.proteinatlas.org/download/pathology.tsv.zip)*(30)*. Genes with prognostic significance were merged by official gene symbol and curated into gene sets.

### Co-expression network analysis

Gene-gene co-expression analysis was performed by computing the Spearman correlation matrix of all genes, as well as a null distribution of correlation coefficients from 10,000 random permutations of the gene expression values. Multiple testing correction was applied using the *qvalue* package in Bioconductor*(57)*. Co-expressed gene pairs were designated as having absolute correlation coefficient ≥ 0.7 and q-value < 0.01. Community detection was performed using Leiden clustering with a resolution parameter of 0.5*(58)*. Visualizations of the resulting correlation network were produced by Gephi using the ForceAtlas 2 layout algorithm*(59)*. Edge weights were set to the correlation coefficient raised to the fourth power. Cluster enrichment analysis was performed using the “enricher” function from the *clusterProfiler* package*(60)*.

## List of Supplementary Materials

Fig S1 to S8

Tables S1 to S11 (included in Data File S1)

Data File S1 (Contains Tables S1 to S11)

## Acknowledgements

The authors thank Hariharan K. Iyer for critical review of statistical and bioinformatics methodology. The authors also thank the Duke University BioRepository & Precision Pathology Center (Duke BRPC; supported by P30CA014236) and the National Cancer Institute’s Cooperative Human Tissue Network (CHTN; supported at Duke University by UM1CA239755) for the provision of samples.

## Author contributions

Conceptualization: PJA

Methodology: MKI, CS, AME, AF, PJA

Investigation: MKI, CS, AME, AF, DPN, PJA

Visualization: MKI, DPN, AME, PJA

Funding acquisition: AME, PJA

Project administration: AF, PJA Supervision: PJA

Writing – original draft: MKI

Writing – review & editing: MKI, DPN, CS, AME, PJA

## Data and code availability

Code and data have been deposited at Zenodo: DOI 10.5281/zenodo.7479047

Code is also available at GitHub:

https://github.com/mkiyer/spatial_rna_ipmn

## Figures

High-resolution images have been uploaded to the manuscript submission website

